# A Metagenomic Study of Antibiotic Resistance Across Diverse Soil Types and Geographical Locations

**DOI:** 10.1101/2024.09.30.615846

**Authors:** Stephanie Pillay, Yasin Tepeli, Paul van Lent, Thomas Abeel

## Abstract

**Background:** Soil naturally harbours antibiotic resistant bacteria and is considered to be a reservoir for antibiotic resistance. The overuse of antibiotics across human, animal, and environmental sectors has intensified this issue leading to an increased acquisition of antibiotic resistant genes by bacteria in soil. Various biogeographical factors, such as soil pH, temperature, and pollutants, play a role in the spread and emergence of antibiotic resistance in soil. In this study, we utilised publicly available metagenomic datasets from four different soil types (rhizosphere, urban, natural, and rural areas) sampled from nine distinct geographic locations to explore the patterns of antibiotic resistance in soils from different regions.

**Results:** *Bradyrhizobium* was predominant in vegetation soil types regardless of soil pH and temperature. ESKAPE pathogen *Pseudomonas aeruginosa* was prevalent in rural soil samples. Antibiotic resistance gene families such as 16s rRNA with mutations conferring resistance to aminoglycoside antibiotics, OXA β-lactamase, ANT(3’’), and the RND and MFS efflux pump gene were identified in all soil types, with their abundances influenced by anthropogenic activities, vegetation, and climate in different geographical locations. Plasmids were more abundant in rural soils and were linked to aminoglycoside resistance. Integrons and integrative elements identified were associated with commonly used and naturally occurring antibiotics, showing similar abundances across different soil types and geographical locations.

**Conclusion:** Antimicrobial resistance in soil may be driven by anthropogenic activities and biogeographical factors, increasing the risk of bacteria developing resistance and leading to higher morbidity and mortality rates in humans and animals.

## Introduction

Antimicrobial resistance (AMR) has contributed to approximately 4.95 million deaths worldwide (Ramtahal *et al*., 2022). The continuous use of antibiotics in clinical and environmental sectors such as animals, soil and water has led to the development of resistant bacteria, which can spread between humans, animals, and the environment (Ramtahal *et al*., 2022). As a result, antibiotic resistance has been identified as a major driver of mortality and one of the most critical environmental issues (Wang *et al*., 2021).

Soil is a large reservoir of microbial diversity and a majority of antibiotics used in clinical and non-clinical sectors have been isolated from soil microorganisms (Wang *et al*., 2021). Interestingly, environmental bacteria in soil possessed antibiotic resistance genes (ARGs) pre-dating the discovery of antibiotics which ensured their survival in the natural environment (Song *et al*,. 2021, Ordine *et al*,. 2023). Despite the natural presence of ARGs in the soil microbiome, anthropogenic activity has impacted the intrinsic resistome. Fertilisation of crops, irrigation, excessive use of xenobiotics in crops, antibiotic use in livestock production and deforestation can alter the microbial community in the soil and disseminate ARGs throughout the environment (Ordine *et al*,. 2023). This makes soil a reservoir for both intrinsic and acquired antibiotic resistance due to the mixture of ARGs from indigenous microbes and those introduced by human activities (Song *et al*,. 2021, Wang *et al*,. 2021).

Biogeographical patterns can also influence antibiotic resistant bacteria and ARGs in the soil (Song *et al*,. 2021). Factors such as pH, temperature, moisture, and nutrients can affect ARG profiles and microbial composition. Soils with a neutral pH, low temperature, high moisture content and nutrient-rich will have an increased abundance and diversity of bacteria as well as a diverse range of ARGs (Wu *et al*,. 2023). These factors promote the spread of antibiotic resistance by mobile genetic elements (MGEs) which enables non-pathogenic and pathogenic bacteria to acquire resistance increasing the risk of bacterial infections in humans and animals (Han *et al*., 2022).

Additionally, the soil microbiome and ARGs can differ across soil types. Natural and pristine soil is largely unaffected or undisturbed by human activity or external sources, serving as a reference point for understanding microbial communities and ARGs with little anthropogenic selection pressure (Van Goethem *et al*., 2018). The rhizosphere soil surrounds the plant roots and is influenced by root secretions and agricultural practices. Urban soil encompasses anthropogenic soil near forests, parks, gardens, and residential areas, while rural soil includes non-urban areas like small villages, towns, and settlements. Soil with more human activity is expected to contain a mixture of environmental and pathogenic bacteria; and resistance genes that confer resistance to clinically relevant antibiotics, heavy metals, biocides and disinfecting agents (Cytryn, 2013). It should be noted that similar soil types from different geographical locations will not necessarily have similar microbial communities or ARGs as agricultural activities, travel, environmental contamination, antibiotic usage, soil management practices, population and socioeconomic conditions differ between countries (Manyi-Loh *et al*., 2018). These factors have complex and interrelated effects on antibiotic resistance in soil.

Since soil encompasses different environmental conditions with distinct microbial environments, it is important to understand how these factors interact and contribute to AMR to develop effective mitigation strategies (Manyi-Loh *et al*., 2018, Wu *et al*,. 2023). In this study, we analysed metagenomic sequences from four different soil types: natural, urban, rhizosphere, and rural soil collected from nine different geographical locations i.e., South Africa, Singapore, China, Israel, Botswana, Chile, Germany, El Salvador, and Antarctica. We aim to determine the differences in microbial composition between each soil type sampled from these various geographical locations. Furthermore, we investigate the differences in antibiotic resistance genes and mobile genetic elements (plasmids, integrons and integrative elements) identified in each soil type with regards to geographical location.

## Methods

### Data sources

Whole metagenomic datasets were obtained from NCBI SRA (https://www.ncbi.nlm.nih.gov/sra). These datasets represented various soil types: natural soil (NS), urban soil (US), rhizosphere soil (RHS) and rural soil (RS) from different geographical locations (South Africa, Singapore, China, Israel, Botswana, Chile, Germany, El Salvador, and Antarctica) sampled over the last 10 years. Only metagenomic sequences that were sequenced using Illumina, had a read base count over 1e+09 and an average read length over 150bp were included. Detailed information about the metagenomes retrieved from the database is included in Supplementary Table 1 and presented in summarized form in Table 1.

**Table 1:** Summary of included studies, respective sample source, location and sample size. Study identifiers start with the soil abbreviation: natural soil (NS), urban soil (US), rhizosphere soil (RHS) and rural soil (RS).

| Study | Sample source | Location | Number of samples | Citation |
| --- | --- | --- | --- | --- |
| NS1 | Deep forest | China | 3 | (Zheng *et al.*, 2021) |
| NS2 | Mountain | China | 5 | (Liao *et al.*, 2022) |
| NS3 | Desert | Israel | 12 | (Bay *et al.*, 2020) |
| NS4 | Desert | Chile | 11 | (Hwang *et al.*, 2021) |
| NS5 | Antarctic | Antarctica | 27 | (Franco *et al.*, 2022) |
| US1 | Forest park,<br>residential areas,<br>hospital areas | China | 10 | (Zheng *et al.*, 2021) |
| US2 | Road verge, park | China | 10 | (Liao *et al.*, 2022) |
| US3 | Nature reserve | Germany | 8 | (Mantri *et al.*, 2021) |
| RHS1 | Greenhouse | Singapore | 13 | (Bandla *et al.*, 2020) |
| RHS2 | Soybean farm | South Africa | 12 | (Babalola *et al.*, 2022) |
| RHS3 | Peanut and maize<br>farm | China | 21 | (Dong *et al.*, 2022) |
| RS1 | Rural village | El Salvador | 25 | (Pehrsson *et al.*, 2016) |
| RS2 | Rural village | Botswana | 1 | (Brooks *et al.*, 2023) |

### Bioinformatic analysis

Quality control of reads was conducted using FastQC v.0.11.9 (Andrews, 2010) and filtered with Trimmomatic v.0.39 (Bolger *et al*,. 2014). Reads were trimmed for quality and adapter contamination using Trimmomatic with the following parameters specified: *ILLUMINACLIP: adapters.fa:2:30:10 LEADING:3 TRAILING:3 MINLEN:30 SLIDINGWINDOW: 4:20*. The microbiome was analysed using Kraken2 v.2.1.0 (Wood *et al*,. 2019) with the minikraken2 microbial database (2019, 8GB). Species confirmation and abundance estimation were performed with Bracken v.2.6.2 (Lu *et al*,. 2017). The relative abundances of bacterial genera were used to determine the microbial populations in each soil type. Statistical significance of the microbiome between studies in a soil type was assessed using the non-parametric Wilcoxon rank sum test and Friedman’s ANOVA with SPSS software (IBM SPSS Statistics). Differences in the microbiome were considered statistically significant at p < 0.05. The Beta diversity of the microbiome among soil types was visualized using non-metric multidimensional scaling (NMDS) analysis with Bray-Curtis dissimilarity in the R package vegan v.2.6-4.

The filtered reads were *de novo* assembled using metaSPAdes v.3.15.2 (Nurk *et al*., 2017) on meta mode with default parameters. The assembled metagenomes were annotated to determine the antibiotic resistance gene families and drug classes using the Comprehensive Antimicrobial Resistance Database Resistance gene identifier (CARD - RGI) v.3.1.2 with default settings including loose matches and the -low_quality -clean options (Jia *et al*., 2016). The version of the database was consistent throughout the analysis. Moreover, the studies in each soil type were compared by analysing the relative abundance of annotated antibiotic resistance genes. This comparison was based on normalizing the data using the total number of datasets in each study indicating their respective proportions.

*De novo* metagenomic assembled contigs were aligned to detect integrons and integrative elements by aligning them to the INTEGRALL database (Moura *et al*,. 2009) and the ICEberg database (Liu *et al*,. 2019). Alignments were done using BWA-mem v.0.7.10 with default parameters (Li & Durbin, 2009) which generated a SAM file. To filter out soft and hard clipped reads and undesirable alignments, the SAM files were processed using the Samclip tool with default parameters to remove alignments that could generate downstream issues. The resulting SAM file was then converted to a BAM file using SAMtools version v.1.9 (Li & Durbin, 2009). Contigs that aligned with databases were considered integrons and integrative conjugative elements. Contigs were classified as plasmid-originating using Plasclass v.0.1.1 (Pellow *et a*, *l.* 2020), with a minimum length threshold of > 1000 and a probability threshold of > 0.75 for contigs to be considered plasmid-originating. To determine if the detected plasmids were linked to antibiotic resistance gene families, the plasmid-originating contigs were compared to previously identified contigs carrying antibiotic resistance genes and matched to corresponding AMR drug classes.

## Results and discussion

### Microbial communities are dependent on their habitat

To identify the bacterial communities within different types of soil from different locations (Supplementary Table 1), we performed taxonomic classification. Soil hosts a diverse range of microbial communities influenced by abiotic and biotic characteristics, microbial abundances, activity, and community composition. Consequently, there is no “typical” soil microbiome, and bacterial relative abundances vary by soil type (Fierer, 2017).

Bacteria in natural soils are influenced by their habitat and environmental conditions (Supplementary Table 2). In Figure 1, *Bradyrhizobium* dominates the deep forest areas (NS1 and NS2), whereas bacteria like *Rubrobacter*, known for its thermophilic traits, are found in desert-like conditions (NS4), as confirmed by Connon *et al*. (2007) Pajares & Bohannan (2016) and Zheng *et al*. (2021). The microbial communities detected were statistically different between each of the natural studies indicating environmental influence (p < 0.05). Climate, pH, temperature and rainfall may all influence the abundances and types of bacteria found soil as discussed by Chase *et al*., (2021).

**Figure 1:**
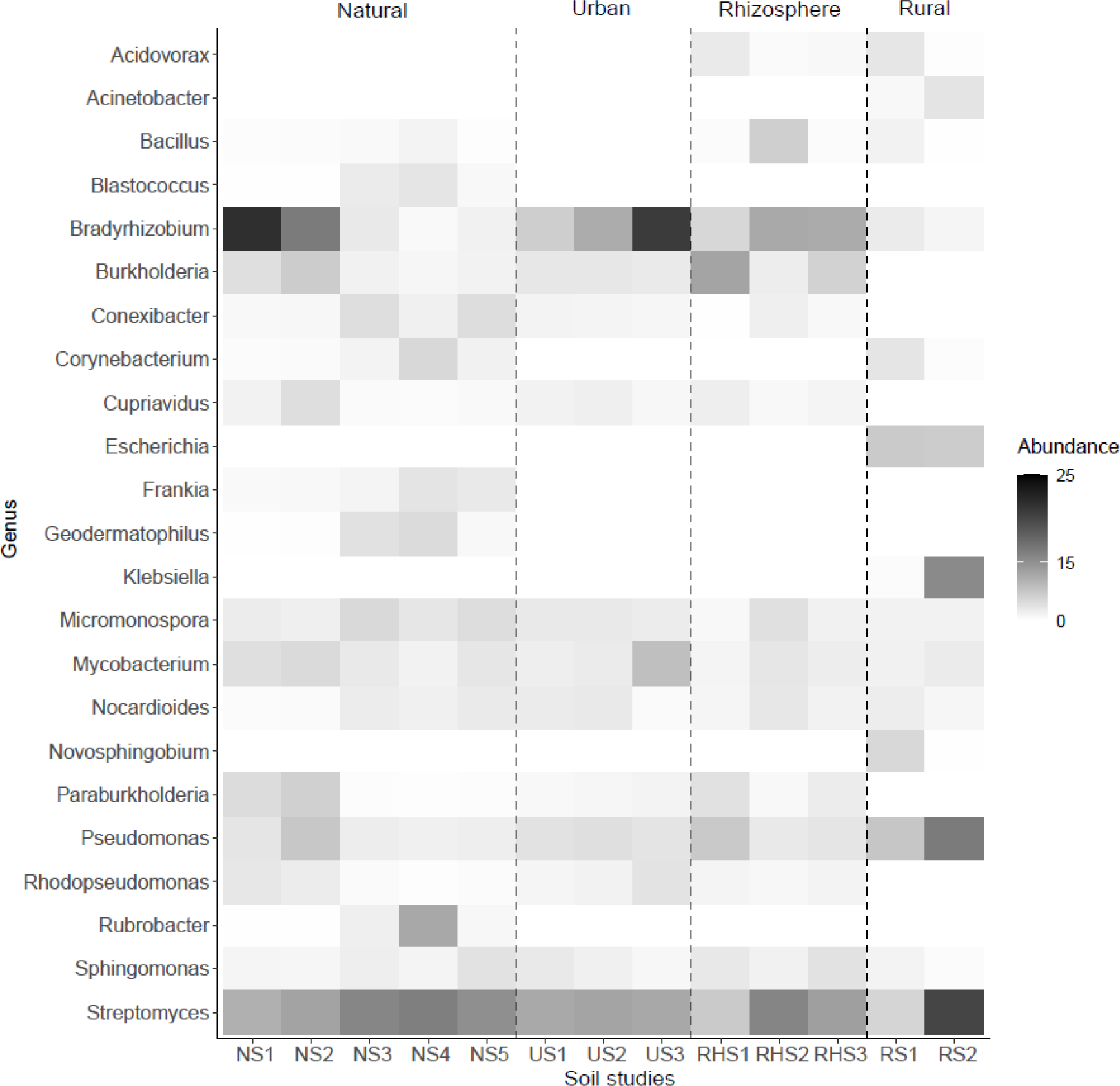
Heatmap showing the relative abundances of bacterial genera over 2% in each study from each soil type. The y-axis indicates the genera, and the x-axis represents the soil studies (Table 1). Each soil type is divided by the dashed black line and the abundances of bacterial genera are indicated by the intensity of grey.

*Streptomyces* was prevalent across all urban studies; however, the overall microbial community differed significantly between these urban studies (p < 0.05) (Figure 1 and Supplementary Table 3). As expected, higher abundances of *Bradyrhizobium*, *Mycobacterium*, and *Rhodopseudomonas* were observed in urban soil obtained from the nature reserve (US3). These bacteria thrive in areas with more vegetation and reduced anthropogenic activity, such as nature reserves, compared to residential and forest park areas (Wong *et al*,. 2014, Gorovtsov *et al*., 2020, Glushakova *et al*., 2021).

Plant growth-promoting bacteria such as *Streptomyces*, *Burkholderia*, *Pseudomonas*, and *Bradyrhizobium* were found in less than 12% abundance across different rhizosphere soil studies (Figure 1 and Supplementary Table 4). Overall, these microbial communities were different in each of the rhizosphere soil studies (p < 0.05). These bacteria also assist in protecting plants against pathogens by producing antibiotics (Kashyap et al., 2019; de Sousa et al., 2022; Liu et al., 2022). *Bradyrhizobium*, which aids in nitrogen fixation and plant growth, is found at an 8% abundance in RHS1, 2% in RHS2 and 4% in RHS3 (Fierer, 2017). Soil in RHS1 and RHS3 consisted of crops supplemented with nitrogen-phosphorus-potassium (NPK) fertiliser (Supplementary Table 1). Despite this, these studies showed a higher abundance of *Bradyrhizobium* compared to RHS2, which was not supplemented. This contradicts previous studies suggesting that *Bradyrhizobium* will be present in a low abundance when nitrogen fertilisers are used (Okubo *et al*., 2013, Yan *et al*., 2017).

Lastly, in rural soil studies (p < 0.05), *Pseudomonas*, *Escherichia*, *Streptomyces*, and *Klebsiella* were prevalent, including the ESKAPE (*Enterococcus faecium*, *Staphylococcus aureus*, *Klebsiella pneumoniae*, *Acinetobacter baumannii*, *Pseudomonas aeruginosa*, and *Enterobacter* species) pathogens *Klebsiella pneumonia* and *Pseudomonas aeruginosa* (Figure 1 and Supplementary Tables 5 and 6 respectively). The high prevalence of these pathogens in rural areas is attributed to anthropogenic factors, such as the presence of domestic animals, open defecation, and inadequate sanitation (Brooks et al., 2023). These conditions contribute to antibiotic resistance hotspots in rural areas, as domestic animals spread faecal matter, and open sewage drains and insufficient sanitation increases the selection pressure for antibiotic resistance (Nadimpalli *et al*,. 2020).

Overall, these results demonstrate that the bacterial composition and abundance within each soil type may be influenced by a myriad of factors, including environmental conditions, nutrients, temperature, oxygen availability, and pH types within the specific geographical location (Pajares & Bohannan, 2016, Fierer, 2017, de Sousa *et al*., 2022). Understanding the composition of microbial communities is critical for comprehending soil health and potential risks associated with antibiotic resistance in various environments.

Natural, urban and rhizosphere soil types have similar microbial communities.

To assess the differences between microbial communities, present in different types of soil from different geographical locations (Table 1), we analysed the beta-diversity. Additionally, we aimed to determine if the geographical locations influence the microbial communities in each soil type. The beta-diversity quantifies the “distance” between microbial communities using the Bray-Curtis index and can be visualized by non-metric multidimensional scaling (NMDS). A smaller distance between samples from different countries and soil types indicates a greater similarity in microbial composition. Figure 2 shows the NMDS plots for each type of soil from their respective countries.

**Figure 2:**
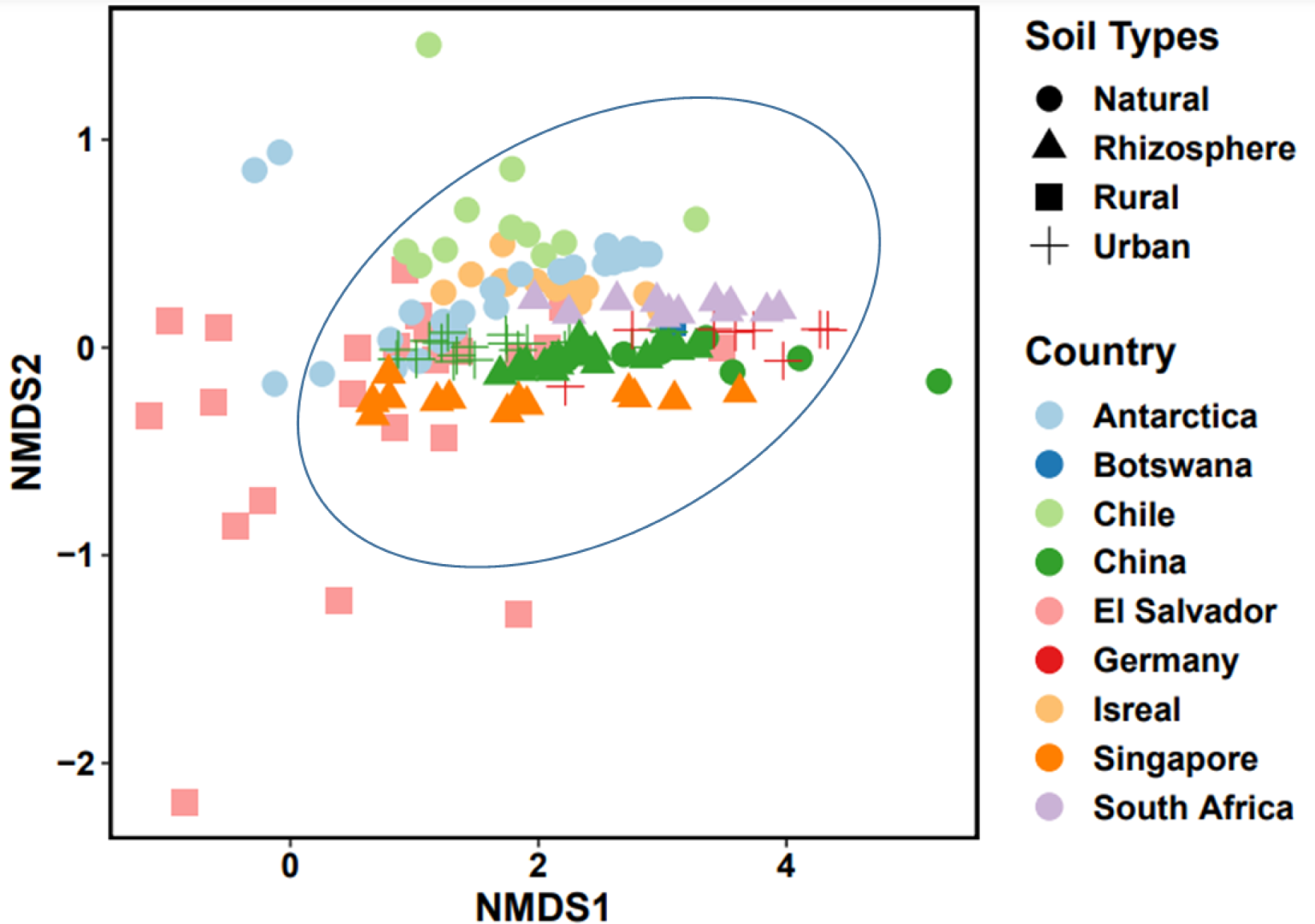
Non-metric multidimensional scaling (NMDS) visualising the dissimilarity (Beta-diversity) between microbial communities from different soil types from different countries (Table 1). Each soil type is represented by a different shape (natural = circle, rhizosphere = triangle, rural = square, urban = cross) and each country or region is colour-coded (Antarctica = light blue, Botswana = dark blue, Chile = light green, China = dark green, El Salvador = pink, Germany = red, Israel = salmon, Singapore = orange, South Africa = lilac). The samples that have smaller distances between them are circled in blue.

Figure 2 shows that most samples cluster together regardless of the soil type or geographical location. Natural, rhizosphere, urban, and many rural soil samples are closely grouped, suggesting similar microbial communities across these soil types. This similarity may be related to vegetation and varying levels of anthropogenic activity (Pajares & Bohannan, 2016, Zheng *et al*,. 2021). However, some rural samples from El Salvador differ which could be due to human activity or soil around latrine toilets and clothes washing areas. This can limit plant-promoting bacteria and support enteric pathogens (Pehrsson *et al*,. 2016, Brooks *et al*,. 2023). Samples from specific studies like RS1 (El Salvador), NS2 (China), NS4 (Chile) and NS5 (Antarctica) are further apart, indicating variations in the microbial composition. Such differences within the same study may result from spatial variability in the soil environment, even when sampling sites are only a few centimetres apart (Fierer, 2017).

Anthropogenic activities, vegetation, and climate play a role in the selection and prevalence of antibiotic resistance genes.

To understand the connection between AMR gene families and the different types of soil, we examined the presence and abundance of AMR gene families in the assembled metagenomes (contigs) from each study within their respective soil type (Table 1). While AMR gene families are naturally present in soil environments, external factors like livestock and human activities can influence their types and abundance (Hernández *et al*., 2023).

Overall, more than 0.001% of contigs contained AMR gene families in each study. We identified five AMR gene families that were abundant (Figure 3). Two of which are associated with acquired resistance: 23S rRNA, which confers resistance to macrolides, and 16S rRNA, which confers resistance to aminoglycosides. Intrinsic resistance gene families include the major facilitator superfamily (MFS) and resistance-nodulation-cell division (RND) antibiotic efflux pumps, and the OXA beta-lactamase gene family, which is associated with acquired and intrinsic resistance (Poirel *et al*., 2010, Garcia *et al*., 2022). These gene families confer resistance to macrolides, and beta-lactams including carbapenems, cephalosporins, and penams (Supplementary Table 7).

**Figure 3:**
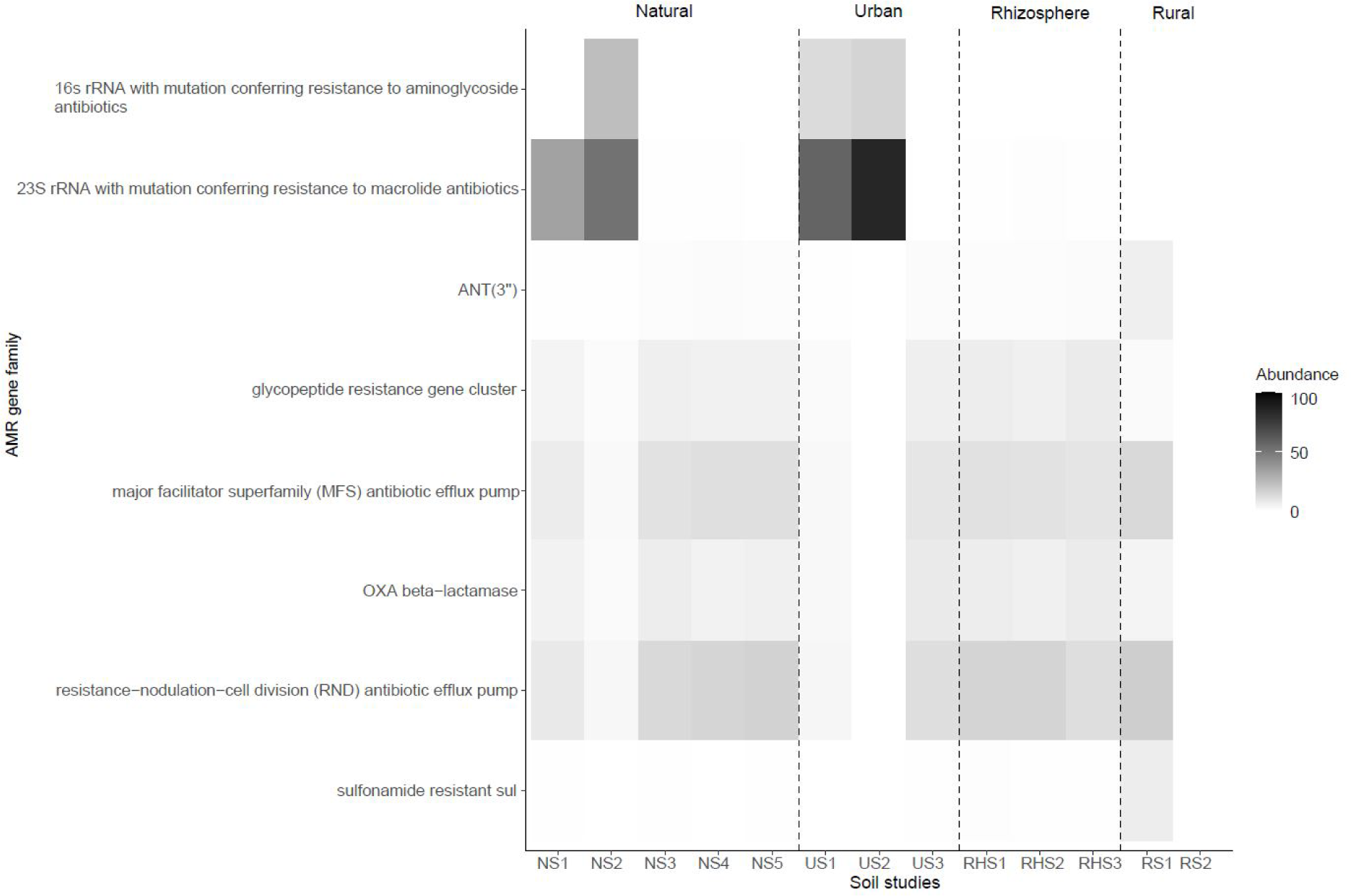
Heatmap showing the abundance of AMR gene families present over 5% in each study within each soil type. The y-axis indicates the AMR gene families, and the x-axis represents the soil studies (Table 1). Each soil type is divided by a dashed black line and the abundances of AMR gene families are indicated by the intensity of grey.

Figure 3 shows that the 23S rRNA mutation conferring resistance to macrolide antibiotics were more prevalent in urban soils as compared to the natural, rhizosphere and rural soils. Specifically, in US2 from China, 85% of the detected AMR gene families were the 23S rRNA gene with this mutation, with other urban studies like US1 had 58% abundance. In contrast, natural soil studies such as NS1 and NS2 had lower abundances of 33% and 52%, respectively. Urban soil samples were collected from areas with high human activity (forest parks, residential zones, and road verges) which have greater antibiotic pollution and selective pressures promoting the proliferation of antibiotic resistant bacteria, as detailed by Osbiston *et al*. (2021).

The MFS and RND efflux pump gene families were detected across all types of soil except for US2 and RS2 (Figure 3). Efflux pumps are key for intrinsic resistance, providing bacteria with a baseline level of resistance by actively expelling a broad range of antibiotics from their cells (Garcia *et al*., 2022). In our study, RS1 exhibited a high abundance of these efflux pump gene families, with 17% for RND and 13% for MFS (Figure 3 and Supplementary Table 7). The presence of these efflux pump gene families in rural soils may also be influenced by factors such as animal feedlots, domestic sewage, and human-excrement-irrigated vegetable fields (Cheng *et al*., 2020). Additionally, vegetation, flora and fauna influences the presence of efflux pump gene families, as discussed by Martinez *et al*. (2009) and Pasqua *et al*. (2019).

Lastly, we found the OXA beta-lactamase gene family present in all rhizosphere and natural soil studies, except US2 and RS2, with a high abundance in US3 (7%) and RHS3 (7%) (Figure 3 and Supplementary Table 7). The OXA beta-lactamase gene family has been shown to confer both acquired and intrinsic resistance and can be found in all soil types; from primitive natural soils with little to no anthropogenic activity to urban environments (Djenadi *et al*., 2018, Han *et al*., 2022). Previous research also suggests that the climate affects the abundance of ARGs, with beta-lactam resistance genes being more common in warmer areas (Li *et al*,. 2022). However, our study found similar AMR gene families across all soil types, regardless of temperature.

### Plasmids linked to aminoglycoside resistance are common in all soil types

To investigate the presence of plasmids and their connection to antibiotic drug classes in different soil types, we analysed the percentage of contigs classified as plasmids in each study and their association with antibiotic drug classes. Although AMR is a natural occurrence in soil, human activity has led to increased contamination with antibiotics and ARGs in the soil. Soil plasmids can transfer these ARGs from one bacterial host to another integrating into plant and animal cells and spreading AMR (Meng *et al*., 2022).

Firstly, less than 1% of contigs were classified as plasmids in each study from every soil type. Further analysis linking detected plasmids to AMR drug classes revealed that, except RHS2, RHS3, RS2 and US2, all studies had more than 0.01% of plasmids associated with AMR drug classes. Notably, RS1 had 1.8% of plasmids linked to AMR. A high percentage of plasmids is common in rural communities due to unregulated access to antibiotics, limited clean water and human and animal activities. These conditions increase the risk of exposure to plasmids and ARGs from effluents, contaminated soils, and waste, enhancing plasmid accumulation and horizontal gene transfer compared to natural and urban soils (Pehrsson *et al*., 2016, Brooks *et al*., 2023).

Secondly, we examined the association between plasmids and specific antibiotic drug classes (Supplementary Table 8). Overall, all studies from different soil types, except RHS2, RHS3, US2 and RS2, contained plasmids associated with aminoglycoside, carbapenem; cephalosporin; penam, glycopeptide and peptide drug classes. Details on the abundances of plasmids linked to AMR drug classes and the individual AMR gene families can be found in Supplementary Table 8.

Plasmids encoding naturally occurring antibiotics were identified in natural and urban soil studies. Among the natural soil types, NS1 showed a high abundance of plasmids associated with aminoglycosides (13%) and carbapenem; cephalosporin; penam (16%) drug classes. These high abundances were consistent with other natural studies (Supplementary Table 8). Similar to NS1, US1 had a high abundance of plasmids associated with aminoglycosides (13%), carbapenem; cephalosporin; penam (25%) along with fluoroquinolones (13%), glycopeptide (25%), macrolide (13%) and tetracycline (13%) drug classes. Additionally, US3 had plasmids linked to similar antibiotic drug classes: aminoglycosides (16%), carbapenem; cephalosporin; penam (11%), glycopeptide (12%) and peptides (9%). In our study, the natural and urban studies, sampled from forests and deserts for natural soils and residential areas and forest parks for urban soils, showed higher abundances of plasmids associated with naturally occurring antibiotics such as aminoglycosides, glycopeptides, and beta-lactams (Zhao *et al*., 2019, Meng *et al*., 2022). These abundances may also be influenced by heavy metals and other contaminants in the soil types as noted by Zhao *et al*. (2019).

Amended rhizosphere soil (RHS1) showed a high abundance of plasmids associated with AMR drug classes. Specifically, aminoglycoside (17%), carbapenem; cephalosporin; penam (8%), glycopeptide (10%) and sulphonamide (10%) resistance plasmids were detected in rhizosphere soils. Although plasmids were detected in RHS2 and RHS3, none were associated with previously identified ARGs. Amended soils, such as rhizosphere soils and natural soils are nutrient-rich environments that can facilitate the spread of AMR through plasmids (van Elsas & Bailey, 2002, Perry & Wright, 2013).

Lastly in rural soils, RS2, a single sample study, did not contain any plasmids associated with AMR. However, RS1 had a high abundance of plasmids linked to aminoglycoside (24%), carbapenem; cephalosporin; penam (9%) and sulphonamide (39%) resistance. This is consistent with a study from a rural village in India that found resistance to carbapenems, cephalosporins, penams, sulphonamides, tetracyclines, and quinolones, with *E. coli* carrying plasmids for resistance to quinolones, cephalosporins, and colistin. This study suggested that the local antibiotic use patterns supports the persistence of plasmids with ARGs (Purohit *et al*., 2017).

While antibiotics and ARGs naturally occur in soil, their transmission to different environments through plasmids is concerning, especially in environments with increased anthropogenic activity. This contributes to the spread of AMR to both pathogenic and non-pathogenic bacteria, posing a significant threat to the natural environment and human health (Meng *et al*., 2022).

### AMR genes associated with integrons that are highly abundant were selected due to the constant use of antibiotics

To identify the presence of integrons and their association with AMR gene families, we classified integrons from the metagenomic contigs present in each soil study. Furthermore, we linked the classified integrons with AMR gene families to determine the differences between soil studies. Integrons are mobile genetic elements that capture and express ARGs, particularly among gram-negative pathogenic bacteria. Integrons are present in all environments and can serve as markers for tracing sources of pollution (Gillings, 2014).

We used the assembled metagenomic contigs to determine the percentage of classified integrons (Supplementary Table 9). In all soil types studied, less than 0.001% of contigs were identified as integrons. The integrons that were found were classified into their respective classes of 1, 2 and 3 (Supplementary Table 9).

Class 1 integrons made up a higher percentage (> 11%) of the integrons we found between all soil types from all studies. Class 3 integrons constitutes a range of 2% to 5% of integrons found while class 2 integrons had a lower percentage of 1% to 2%. The remainder of the integrons were unclassified. These high abundances of class 1 and class 3 in all types of soil regardless of country are expected as class 1 and 3 integrons are found in *Proteobacteria* in soil environments whereas class 2 integrons are commonly found in marine environments. Overall, integrons are found in diverse environments which include forest soil, desert soil, Antarctic soil and plant surfaces (Gillings, 2014).

In our study, we identified class 1, 2 and 3 integrons and their association with AMR genes (Supplementary Tables 10 and 11). We found a high abundance of class 1 integrons associated with disinfecting agents and sulphonamide resistance in all types of soil. The high abundance of the *qacE* gene with class 1 integrons can be traced back to the use of quaternary ammonium compounds which were first used as hospital disinfecting agents in 1930 and have become a part of the class 1 integron, known to be highly abundant in soils (Gillings, 2014, Cheng *et al*., 2020). Similarly, sulphonamides were the first true antibiotic to be used in the 1930s and have been selected for and can be commonly found in class 1 integrons (Gillings, 2014).

A high abundance of class 2 integrons was associated with aminoglycoside resistance (*aadB*) which ranged from 15% in the RS1 to 75% in NS3 and NS4, and 83% in RHS3 and US1 (Supplementary Tables 10 and 11). Aminoglycoside antibiotics were originally isolated from soil bacteria and commonly used in agriculture, increasing their presence in soil environments, potentially affecting plant-soil microbial communities. They also enter soils through human waste, municipal wastewater systems, and clinical use. These direct and indirect exposures contributes to the detection of aminoglycoside resistance genes in various soil types, often associated with class 1 and class 2 integrons (Blau *et al*,. 2018, Coates *et al*,. 2022). Previous studies have shown that the *aadB* aminoglycoside resistance gene is frequently associated with class 1, 2, and 3 integrons and is present in *Pseudomonas aeruginosa*, *Salmonella* spp., *Acinetobacter baumannii*, *Klebsiella* spp., and *Escherichia coli* which are present in our study (Sabbagh *et al*., 2021). Additionally, Jones *et al*., (2005) showed that the *aadB* gene, associated with class 1 integrons, had a close association with the *bla*SHV gene conferring resistance to penicillin and cephalosporins. This association was not seen in our study. Lastly, class 3 integrons were predominantly associated with the *qacE* gene, which provides resistance to disinfectants. Over 55% of class 3 integrons in all soil studies were linked to the *qacE* gene, with 100% in NS4, 97% in RHS2, and 97% in US3. Similarly to class 1 integrons, the *qacE* gene are commonly found in class 3 integrons as it is considered to be a conserved segment and had been selected for in the past (Sabbagh *et al*., 2021).

### Integrative and mobilizable elements were identified in each type of soil

Finally, we used the assembled metagenomic contigs to determine the percentage of integrative elements. Integrative elements consist of integrative and conjugative elements (ICE), integrative and mobilizable elements (IME) and cis-integrative and mobilizable elements (CIME) that are self-transmittable and can carry and spread ARGs to other bacterial hosts (Delavat *et al*., 2017).

We found that all the studies across all soil types had less than 0.1% of contigs classified as integrative elements. ICE elements were the most prevalent integrative element compared to IMEs and CIMEs. In the RS2 study, 100% of the integrative elements were CIMEs, while other studies had less than 20% CIMEs. The abundance of ICEs in all studies across all soil types was over 70%, with no ICE elements found in RS2.

The most prominent ICE families among all studies in all soil types were *Tn4371* (>12%), *SXT/R391* (>5%) and *ICEclc* (>8%) (Supplementary Table 12). These ICE families are commonly found in soil and aid in the degradation of pollutants (Obi *et al*., 2018, Hirose *et al*., 2021, Roman *et al*., 2021). Additionally, we identified different SGI1 IME families present in the different studies within the different types of soil (Supplementary

Table 13). SGI1 has been characterized as a *Salmonella* genomic island that can spread multi-drug resistance to human and animal pathogens (Cummins *et al*,. 2020). Bacteria carrying these integrative elements have a higher potential for spreading infections thus facilitating the spread of AMR to other environments (Delavat *et al*., 2017). This underscores the importance of continuously monitoring mobile genetic elements in different soil types.

## Conclusion

We analysed publicly available metagenomic data to investigate the microbial population, antibiotic resistance genes and mobile genetic elements in rhizosphere, urban, natural, and rural soils sampled from South Africa, Singapore, China, Israel, Botswana, Chile, Germany, El Salvador, and Antarctica to understand the spread of antibiotic resistance in different soil environments.

Our study revealed that plant-promoting bacteria such as *Bradyrhizobium*, dominates soil rich in vegetation and nutrients such as the rhizosphere and natural soil regardless of geographic location. Urban soils contain a mix of plant-promoting bacteria and pollutant-degrading bacteria, such as *Arthrobacter*, while rural village soils harbour a combination of environmental bacteria and opportunistic pathogens like *E. coli.* Bacteria in the soil are dependent on physiological (temperature, mixing of soil profiles, water change, soil compaction), chemical (heavy metal, soil pH, polyaromatic hydrocarbons) and biological (artificial vegetation, faecal contamination) factors. As such, more pathogenic bacteria especially those on the ESKAPE pathogen list are dominant in areas with high human activity. These bacteria can carry antibiotic resistant genes that may spread to domestic and food animals, vegetation, water sources and humans leading to untreatable infections in all sectors.

Antibiotic resistant gene families detected in this study i.e., 23S rRNA with mutation conferring resistance to macrolide antibiotics, major facilitator superfamily (MFS) antibiotic efflux pump, resistance-nodulation-cell division (RND) antibiotic efflux pump, OXA beta-lactamase, ANT(3’’) and 16s rRNA with mutation conferring resistance to aminoglycoside antibiotics in soil. These genes favour the development of multi-drug resistance, posing a threat to antibiotic effectiveness. While antibiotic resistant gene families such as the RND and MFS efflux pump gene families occur naturally in soil, anthropogenic activity, vegetation, and climate may play a role in the emergence of AMR and the acquisition of ARGs. Additionally, plasmids associated with AMR gene families like OXA beta-lactamase facilitates the spread of resistance from soil to various environments like water, humans or animals.

Integrons detected in this study were linked to commonly used antibiotics in all soil types, suggesting that the overuse of antibiotics contributes to the spread and emergence of antibiotic resistant bacteria. These bacteria may also possess adaptive strategies provided by integrative elements, *SXT/R391, Tn4371, ICEclc* and IMEs which can potentially carry accessory genes i.e., ARGs. Integrative elements allow bacteria in the soil to stabilise and adapt to environmental conditions and human activity as well as favouring gene transfer and the spread of AMR.

To fully understand the emergence and spread of AMR in the environment, factors such as geographical location, biogeographical patterns, and soil type must be considered. By analysing MGEs and ARGs in diverse soil types from different countries, we can gain insights into how AMR spreads and adapt strategies to manage it effectively. Future research should focus on collecting more rural soil data and considering additional factors such as antibiotic usage, residues, industrial practices, and local regulations to address AMR and environmental pollution.

## Declarations

### Authors’ Contributions

SP and TA designed experiments. SP conducted all experiments, analysed data and wrote the manuscript. YIT and PL assisted in data analysis. SP, YIT, PL and TA proof-read and edited the manuscript.

